# Lifespan heterogeneity reflects intrinsic quality, not trade-offs

**DOI:** 10.1101/2025.09.27.678976

**Authors:** Silvia Cattelan, Dario Riccardo Valenzano

## Abstract

Individuals of the same species often display extensive lifespan heterogeneity. However, the factors underlying such heterogeneity remain largely elusive. Here, we examined whether lifespan variation in the naturally short-lived turquoise killifish (*Nothobranchius furzeri*) is caused by reproductive trade-offs. We found that lifespan and reproduction do not trade off: early-life reproductive success positively predicts male lifespan and reflects intrinsic quality. A causal-informed model, where intrinsic quality affects both survival and reproduction, fully recapitulates the empirical data. Together, our results show inter-individual variation in intrinsic quality explains lifespan heterogeneity in a short-lived vertebrate, challenging the view that reproductive costs drive inter-individual lifespan variation.

## Introduction

Heterogeneity in lifespan among individuals of the same species has been attributed to several causes, including survival–reproduction trade-offs, stochastic variation in early-life exposures, and inter-individual differences in intrinsic quality. Classical life-history theory, grounded in the assumption of limited resources, posits that organisms strategically allocate resources between reproduction and somatic maintenance, thereby generating much of the observed variation in lifespan both within and between species^1-3^. While stochastic processes undoubtedly contribute to individual life histories, work in genetically identical systems indicates that early-life variability is not expressed as irreducible noise but can instead be amplified and stabilized into persistent physiological states that are predictive of lifespan. In isogenic *Caenorhabditis elegans* populations, stochastic differences in the early expression of stress- and aging-associated genes predict remaining lifespan, revealing stable inter-individual differences that emerge despite uniform genetic and environmental conditions^4^. Similarly, recent single-organism transcriptomic analyses demonstrate that early transcriptional heterogeneity can become canalized into coherent gene-expression programs that persist across the lifespan and strongly predict survival outcomes^5^. Together, these findings suggest that stochasticity primarily acts by shaping durable differences in intrinsic individual state, rather than constituting an alternative, age-dependent explanation for lifespan variation. This reframes a central and largely unresolved question: to what extent does heterogeneity in lifespan reflect differences in resource allocation strategies versus persistent differences in intrinsic individual quality.

Sex-specific life-history strategies provide a powerful context in which to address this question. Under classical life-history models, elevated male investment in early-life reproduction is expected to incur costs to somatic maintenance, generating a negative association between reproduction and survival and favouring a “live fast–die young” strategy^6,7^. However, theoretical and comparative studies suggest that sexual competition may instead impose strong selection on overall individual quality^8,9^, challenging the assumption that reproduction and survival are necessarily negatively coupled. In this framework, high-quality males may both reproduce more and survive longer, leading to positive covariation between reproductive output and lifespan^10-12^. Despite its central importance for understanding the origins of lifespan heterogeneity, empirical evidence testing these alternative predictions in males remains scarce (reviewed in^13^).

The turquoise killifish (*Nothobranchius furzeri*) has evolved in ephemeral habitats where individuals can mate within a narrow time window, defined by seasonal water availability. Males show a strong and rapid decline in reproductive functions^14,15^ despite their extremely short lifespan of 4-8 months^16^. Alternative life-history strategies among males may explain both the considerable variation in lifespan among individuals^16-18^ and reproductive aging^14,15^, through the selective disappearance of "live fast−die young" individuals from the population.

To investigate whether the variation in turquoise killifish lifespan reflects differential investment in reproduction among males, we employed a longitudinal experimental design in which we measured male reproductive success after controlled *in vitro* fertilizations (IVF) at three different time points during lifespan and recorded the age of death. Under the trade-off model, we expect to find a negative association between survival and reproductive success, especially in early life. Alternatively, in the absence of obvious trade-offs, individual quality may affect both individual lifespan and reproduction, resulting in positive covariation between reproductive success and lifespan.

To investigate whether the variation in turquoise killifish lifespan reflects differential investment in reproduction among individuals, we first performed IVFs at three different ages during male lifespan: at young (9 wph, 100% population alive), middle (15 wph, 85% population alive) and old (20 wph, 60% population alive) age (**Figure 1a**) by keeping female age constant throughout the experiment. As a measure of reproductive success, we counted the number of fertilized eggs (i.e., fertilization rate) and the number of embryos that survived after 72 hours after fertilization (i.e., embryo survival). Here, we present the results for embryo survival since fertilization rate largely mirrors the patterns observed in embryo survival (see **Supplementary Results, Figure S1**). First, confirming previous work^14^, we found that age significantly affected embryo survival (*X*^2^=26.805, *p*=1.512e-06). Old males produced embryos with significant lower survival compared to young males (*z*=4.572, *p*<0.0001) (**Figure 1b**). Moreover, we found that embryo survival significantly declines already at mid-age (*z*=3.764, *p*=0.0005) (**Figure 1b**). Second, to test for reproduction-survival trade-offs, we estimated posterior predictive distributions of the relationship between lifespan and embryo survival at the three different ages. We fitted Bayesian binomial models^19,20^ with a logit link function modelling the log-odds of embryo survival (i.e., the logit of the embryo survival probability) as a linear function of log transformed lifespan (see *Equation 1*). Contrary to the trade-off expectation, we found no negative correlation between individual reproductive success and lifespan (**Figure 1b**). Moreover, posterior estimates indicate that individuals with higher reproductive success at young age live longer, i.e., early-life reproductive success positively covaries with individual age at death (β_mean_=1.07; 0.51-1.67, 95% credible intervals), while there is no correlation between individual survival and reproductive outcome when sampling fathers at mid (β_mean_=-0.15; - 0.95-0.65, 95% credible intervals) and old age (β_mean_=-0.14; -2.98-2.33, 95% credible intervals) (**Figure 1b**). These results indicate a positive association between embryo survival and lifespan at young age but not later in life.

**Figure 1.**
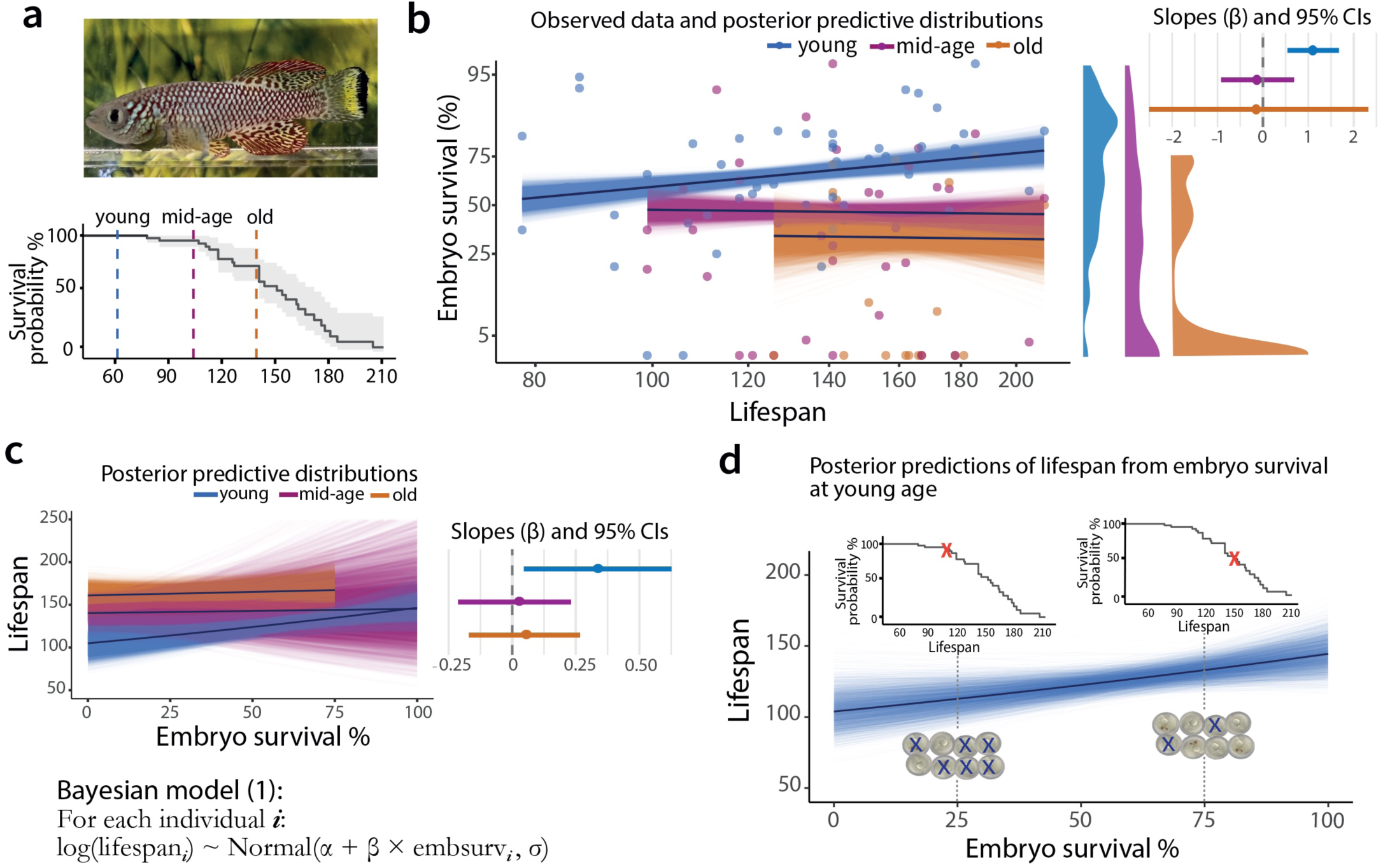
Early life reproductive success predicts individual lifespan. **a)** Typical adult male from the GRZ-D strain and survival curves of the male cohort used in the study (median lifespan: 154 days). At three age classes (young, mid-age and old) across males’ lifespan, we collected the ejaculate and performed IVFs. We measured embryo survival, i.e. proportion of survived embryos at 72 hours after IVF. Lifespan of each male was also recorded. **b)** On the left, scatterplot showing the relationship between lifespan and embryo survival across the three age classes. Dots represent observed data from each individual male. Mean posterior predictions (black lines) and posterior predictive distributions (coloured lines) from Bayesian models are fitted. Embryo survival was modelled as a binomial outcome with a logit link and log-transformed lifespan was added as a predictor in the model. For visualization purposes, observed data for embryo survival and lifespan is transformed using the logit function and the log scale, respectively. In the center, Kernel density plot depicting embryo survival at young (mean±sd: 63.2±24.3%), mid (mean±sd: 40.4±29.6%) and old (mean±sd: 26.6±31.0%) age. Pairwise comparisons are significant (*p*<0.001) for all groups except between mid and old age. On the right, Forest plot of slopes (β estimates) for each age class. Points indicate posterior means and horizontal bars represent 95% credible intervals, relative to zero as the null effect. **c)** Mean posterior predictions (black lines) and posterior predictive distributions (coloured lines) from Bayesian models run for each age class. Lifespan was modelled as the outcome variable. For visualization purposes, lifespan was back-transformed from the log scale by exponentiation. The Equation used to run Bayesian models predicting father lifespan from embryo survival is reported on the bottom. Log-transformed lifespan was modelled as a linear function of embryo survival, where α represents the intercept, β is the slope (i.e. effect of embryo survival on lifespan), and σ is the residual standard deviation. Each model (one per age class) was run with 4 chains, each consisting of 2000 iterations, of which 1000 for warmup. On the right, forest plot of slopes (β estimates) for each age class. Points indicate posterior means and horizontal bars represent 95% credible intervals, relative to zero as the null effect. At young age the posterior distribution of the slope indicates a 95% probability that higher embryo survival is associated with increased male lifespan, as the 95% credible intervals does not include 0. **d)** Example of posterior predictions for father lifespan at young age. To estimate the expected father lifespan for a given embryo survival probability (*p*), we extracted the posterior means of the intercept (α) and slope (β) from the model and calculated the expected lifespan at *p* as follows: Lifespan_(*p*)_ = exp(α_mean_ + β_mean_ × *p*). An increase of 50% in embryo survival (e.g. from 25% to 75%) translates into a 17% increase in lifespan. Lifespan differences are indicated on the survival curves.

Since father survival and embryo survival covary for measures taken at young father age, we asked whether embryo survival statistically predicts variation in father lifespan across individuals. Running a Bayesian model for each age class with father lifespan as the outcome variable (**Figure 1c**), we found, in line with the previous model, that the effect of early-life reproductive success on lifespan is positive (**Figure 1c**). Specifically, the posterior predictive distribution of the slope (young age: β_mean_=0.33; 0.035-0.620, 95% credible intervals, **Figure 1c**) indicates a 95% probability that higher embryo survival is associated with increased male lifespan. To estimate the expected father lifespan for a given embryo survival probability (*p*), we extracted the posterior means of the intercept (α) and slope (β) from the model and calculated the expected lifespan at *p* as follows: Lifespan_(*p*)_ = exp(α_mean_ + β_mean_ × *p*). For instance, an increase of 50% in embryo survival predicts a 17% proportional increase in expected lifespan (**Figure 1d**). When running a model using reproductive success measured at older ages, the credible intervals associated to the slope of embryo survival on lifespan include zero both at mid (β_mean_=0.017; -0.202-0.229, 95% credible intervals) and old age (β_mean_=0.052; -0.164-0.269, 95% credible intervals), indicating that measuring embryo survival from mid and old age fathers provides no information about their lifespan (**Figure 1c**).

Since trade-off models don’t explain the positive covariation between survival and reproduction at the species and individual level, we explored an alternative model, which we formalized as a direct acyclic graph (DAG) (**Figure 2a**), a probabilistic approach that helps define causal relationships among variables, identifying potential confounders, mediators and colliders. We modelled that individual quality (hereafter, *intrinsic quality*) causally impacts both individual lifespan and functional quality. Functional quality affects the ability to reproduce and thus, reproductive success (i.e., embryo survival). Age, in turn, impacts reproductive success both directly and through functional quality (**Figure 2a**). This simple model leads to three important consequences. First, variation in *intrinsic quality* among individuals leads to the positive correlation between survival and reproduction. Second, age negatively impacts embryo survival – but not lifespan! – hence leading to the loss of covariation between reproductive success and lifespan at older age. Furthermore, individuals with low *intrinsic quality* die at young age, reducing the remaining variation of *intrinsic quality* at the population level in middle and old age. Finally, since early-life reproduction is a *bona fide* proxy for *intrinsic quality*, it can be reliably used to predict both parental lifespan and, since it captures both survival and reproduction, Darwinian fitness. Similar to our observed data, the model specified in the DAG showed no negative correlation between embryo survival and lifespan (**Figure 2b**). Moreover, simulated individuals with higher embryo survival at young age live longer, while embryo survival at mid and old age does not correlate with lifespan (**Figure 2b**), meaning that longer-lived individuals have higher early-life embryo survival compared to individuals living shorter (**Figure 2c**). Since the simulated data produced from our generative model recapitulates our observed data, we asked whether embryo survival statistically predicts variation in father lifespan across simulated individuals in an age-dependent manner. Running a Bayesian model for each age class on the simulated data, we found, in line to the observed data (see **Figure 1c**), that the effect of embryo survival on lifespan is positive at young age (**Figure 2d**). Specifically, the credible intervals associated to the slope of embryo survival on lifespan include zero at young age, but neither at mid nor old age (**Figure 2d**). Consistent with our observed data, our generative model confirmed that the positive correlation between reproduction and lifespan is lost at older ages. Overall, these results indicate that *i)* inter-variation in *intrinsic quality* is sufficient to generate the positive correlation between reproduction and lifespan, and *ii)* age weakens the correlation between reproduction and lifespan at older ages.

**Figure 2.**
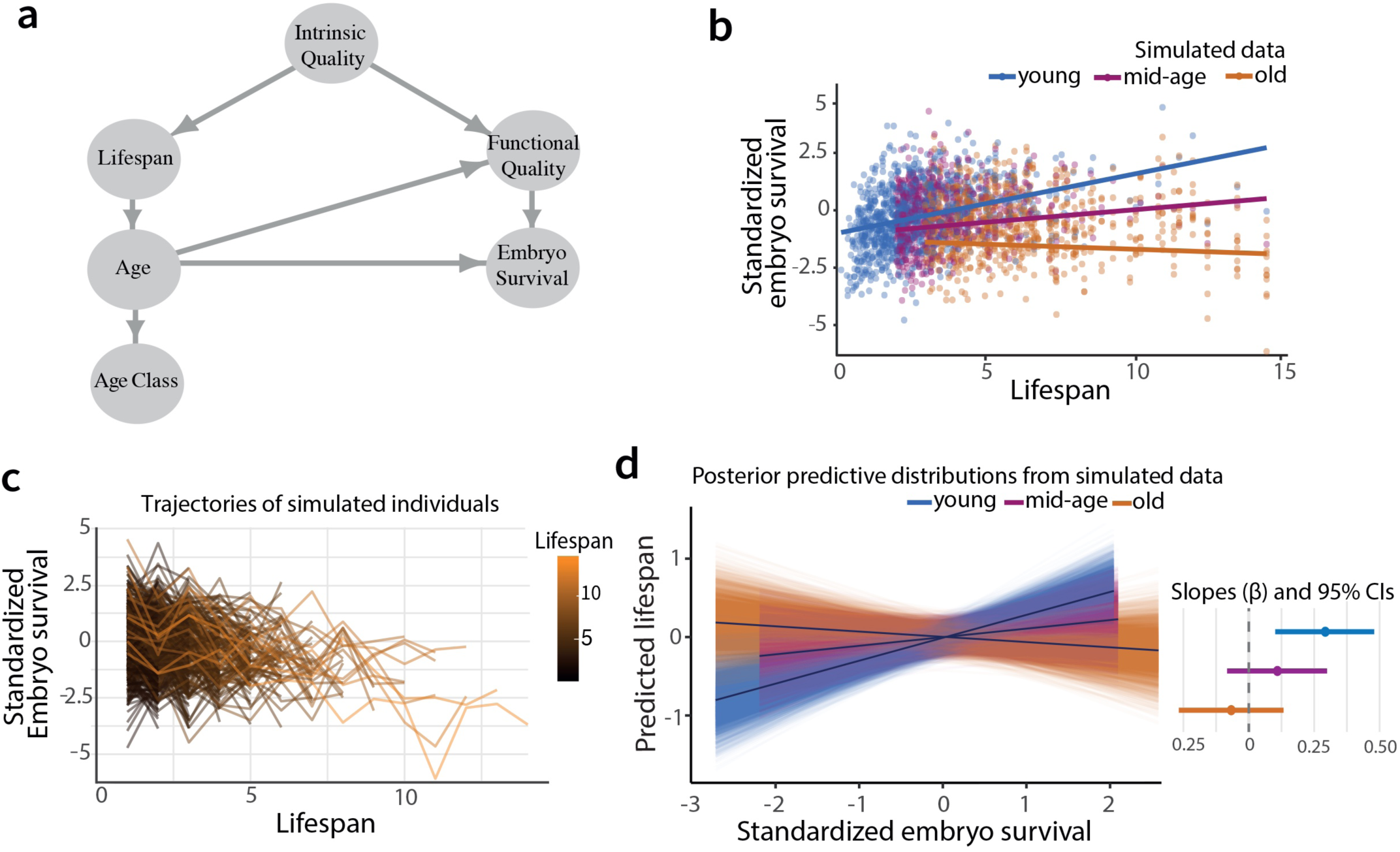
Intrinsic quality explains the age-dependent change in the correlation between individual reproductive success and lifespan. **a)** Direct acyclic graph showing the causal relationships among variables. *Intrinsic quality* influences both lifespan and functional quality, which in turn affects reproductive success (e.g. embryo survival). Age impacts reproductive success both directly and indirectly through functional quality. Longer-lived individuals reach later ages compared to short-lived individuals, making lifespan collinear with age class (e.g. young, mid-age, old). **b)** Scatterplot showing the correlation between embryo survival and lifespan in a simulated population in which individuals differ for their *intrinsic quality*. Lifespan and embryo survival were expressed as linear functions of *intrinsic quality* and age classes were defined according to experimental data thresholds. Each dot represents an individual. Similarly to observed data, simulated individuals with higher embryo survival at young age live longer, while embryo survival at mid and old age does not correlate with lifespan. **c)** Individual trajectories of the relationship between embryo survival and lifespan across ages from simulated data. Each line represents an individual. Individuals with high embryo survival at young age tend to live longer (orange lines) compared to individual with low embryo survival at young age (black lines). **d)** Mean posterior predictions from Bayesian models on the simulated data for each age class. Consistent to observed data, embryo survival at young age but not at mid and old age, predicts individual lifespan, in accordance with the observed data. On the right, forest plot of slopes (β estimates) for each age class. Points indicate posterior means and horizontal bars represent 95% credible intervals, relative to zero as the null effect.

Our longitudinal experimental design revealed that males with higher early-life reproductive success also lived longer than those with lower early-life reproductive success. The absence of a clear trade-off between reproduction and survival at the individual level also rules out the possibility that reproductive aging in killifish arises as a statistical artefact of the selective disappearance of short-lived, high-investing males. These results align with emerging evidence from females of several species, where positive associations between early-life reproduction and survival – rather than trade-offs – define the primary axis of individual variation^15,21^.

We propose that the positive covariation between reproduction and lifespan reflects intrinsic differences in individual quality. High-quality individuals can simultaneously achieve higher reproductive success and extended survival. The variation in *intrinsic quality* among males is sufficient to explain the loss of correlation between reproduction and lifespan at older ages: as low-quality individuals die earlier, the remaining population becomes increasingly homogeneous in quality. Simulations based on our direct acyclic graph (DAG) model recapitulated the main empirical findings, suggesting that *intrinsic quality* provides a common source of heterogeneity shaping both lifespan and embryo survival.

More broadly, our results highlight how life-history trade-offs, while useful and intuitive^22^, might not fully explain life-history relationships observed within or between populations. Our results support an expanded framework of life-history evolution that integrates variation in individual quality and life stages^23,24^. By integrating reproductive performance and survival within a unified quality-based model, our results offer a revised perspective of how reproduction and ageing covary, with implications for understanding the evolution of aging and life-history diversity.

## Materials and Methods

### Fish husbandry

The turquoise killifish used in this study were laboratory-reared fish from GRZ-D strain maintained at the Fish Facility of the Leibniz Institute on Aging (Jena, Germany) under standard laboratory conditions (12:12 light–dark cycle; water temperature 26 ± 1 °C; pH 6.5-7.5). Fish were maintained in 3.2L tanks (density1.6L/fish, Aqua Schwarz recirculating system) provided with enrichments. Fish were fed *ad libitum* twice per day with live brine shrimps (*Artemia salina*) until week 10. From week 4 fish are fed also with bloodworm larvae (*Chironomus spp.*) once per day throughout their life. Extending details on housing and breeding conditions are described in a previous publication^25^. The experiment was carried out under permission of the following animal experimental license: FLI-22-107.

### In vitro fertilizations

To investigate how investment in reproduction affects lifespan, we performed in vitro fertilizations (IVFs) at three different time points throughout male’s lifespan. The average age of males was 62.8±6.2 days (mean±sd) at young age, 103.5±6.8 days at mid-age, and 140±4.7 at old age. Between young and mid-age n=8 (∼16%) males died, while between mid and old age n=20 (∼50% of the males alive until that point) males died. The males that survived until old age were n=20 (∼41% of the initial males). The median lifespan of the males was 154 days (141-176 95% confidence intervals). Before IVF, we standardized recent social/ sexual history by placing each experimental male with two females (age ranging between 80 and 110 days) in a 10L-tank (Acqua Schwarz) and we let them mate for 5 days. To ensure the replenishment of sperm reserves after the mating, each male was isolated from females for 5 days. The male had visual and olfactory access to a female without the possibility to mate with it. At day 10, we performed IVFs by following the protocol that we published previously^26^. To collect sperm from the males, each male was anesthetized in a water bath containing Tricaine (200mg/L) and then placed on a moist sponge under a stereomicroscope (Leica S9D). After drying the urogenital area with a tissue, sperm were collected using a 5-μl microcapillary by gently pressing the male’s abdomen. The ejaculate was then placed in 35μl Hank’s buffer solution and maintained in ice and used within 20 minutes from collection. To collect eggs from the females, each female was anesthetized as above, then dried with a tissue and placed on standard microscope slide. Eggs were collected by pressing the female’s abdomen with an angle chisel sharper. Eggs were then placed in a petri dish and immerged in a 4% BSA solution at room temperature until use. For each IVF trial, we pooled eggs from at least 3-4 females to limit the effect of female quality on fertilization rate and embryo survival (mean±sd number of eggs/IVF trial: 33.1±9.16). Immediately before the IVF, we removed the BSA with a disposable Pasteur pipette and 30μl of the sperm solution was mixed with 120μl of tank water and then added to the eggs. The remaining 5μL of the sperm solution was used to measure sperm concentration using a cell counter (TC20, BioRad). After 5 minutes from the IVF, eggs were immerged in a diluted solution of methylene blue (1:1000) and incubated at 28°C. We performed a total of 108 IVF trials (n=48 at young age, n=40 at mid-age, n=20 at old age) using 72 females in total. The age of the females was maintained constant across the experiment (mean±sd female age: 98.2±7.60 days). The same female was used for a maximum of two IVFs with 2-week break between the trials.

### Fertilization rate and embryo survival

After 3.5 hours post IVF, we checked the fertilization rate by counting the fertilized and unfertilized eggs under a stereomicroscope. Fertilized eggs are scored by the presence of diving cells (two- to four-cell stage). Unfertilized eggs were removed, and the methylene blue solution was changed. The survival of the embryos was then scored 72 hours post fertilization (hpf) by counting the number of embryos still alive. We scored embryo survival at 72hpf, which corresponds to the end of the epiboly stage^27^, because our previous study revealed that 85% of total embryo mortality occurs within this time frame^14^. In an independent pilot experiment, we found a strong positive correlation (*r*=0.50, *p*<0.001) between embryo survival following IVFs and natural matings, suggesting that IVF-derived embryo mortality is not an artefact of the procedure but reflects a biological effect.

### Effect of male age on reproductive success and correlation with lifespan

We ran the following analyses using R v 4.2.0^28^. To test the effect of male age on embryo survival, we fitted a generalized mixed effect model using the “glmer” function in the “lme4” package^29^ and assumed a binomial error distribution and a logit link function. We fitted the model with Age class (levels: young, mid-age, old) as a categorical fixed effect, sperm concentration as a continuous fixed effect (i.e. covariate) and male ID as random effect to account for the non-independence of the data. The model was checked for overdispersion by using the “dispersion_glmer” function within the “lme4” package and corrected by including an observation-level random effect. We calculated *p* values of the fixed effects by Type II Wald chi-square tests using the “Anova” function in the “car” package^30^. To calculate pairwise effects among the different age classes (young *vs* mid-age; mid-age *vs* old; young *vs* old) we performed post-hoc analyses of contrasts with the “lsmeans” function in the “emmeans” package ^31^ using the Tukey method adjusted for multiple comparisons. To investigate the correlation between reproductive success and lifespan we ran Pearson’s correlations between the proportion of embryos alive (or the proportion of fertilized eggs) and lifespan (in days) for each age class and for the all dataset, using the “cor.test” function in the “psych” package^32^. The same statistical tests were also applied to fertilization rate (**Supplementary Results**).

### Bayesian models on observed data

To estimate the parameters that fit the observed relationship between lifespan and reproductive success, we employed a Bayesian approach in Stan^20^ via the R interface “rstanarm”^19^. For each age class (levels: young, mid-age, old), we ran a Bayesian model to assess whether embryo survival at a specific age was associated with male lifespan. We modelled the number of fertilized eggs that survived 72hpf as a binomial outcome with a logit link function. The log-odds of embryo survival were modelled as a linear function of the log-transformed lifespan of the father. The model included an intercept (α) and a slope (β) with weakly informative priors (see *Equation 1*). Each model was run with 4 chains, each consisting of 2000 iterations, of which 1000 for warmup.

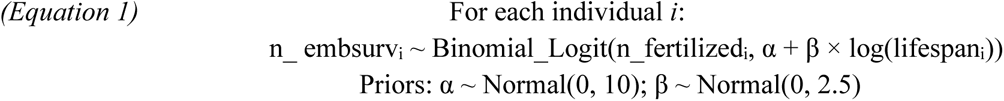

Where: n_embsurv*ᵢ*= number of embryos survived for individual *i;* n_fertilized*_i_*= number of fertilized eggs for individual *i*; α= intercept; β= slope (effect of log-lifespan on embryo survival); and log(lifespan*ᵢ*)= log-transformed lifespan of individual *i.* We performed similar models for fertilization rate by modelling the number of eggs that were fertilized as the binomial outcome. We assessed model convergence using visual inspection of trace plots (“traceplot” function) and checked whether all parameters had R-hat values<1.01, indicating good convergence. Posterior probabilities of the intercept and slope were extracted (“extract” function) for each age class, and the posterior predictions of embryo survival were computed across a regular sequence of 100 father log-lifespan values. We plotted individual posterior predictions and the mean posterior prediction for each age class using “ggplot2” package ^33^ (see associated code).

To estimate the predictive power of embryo survival for father lifespan we ran a Bayesian model for each age class with lifespan as the outcome variable. In these models the log-transformed lifespan was modelled as a linear function of the proportion of embryos that survived 72hpf. The model assumed normally distributed residuals and included an intercept (α), a slope (β), and a residual standard deviation (σ) and weakly informative priors (see *Equation 2*). As before, each model was run with 4 chains, each consisting of 2000 iterations, of which 1000 for warmup.

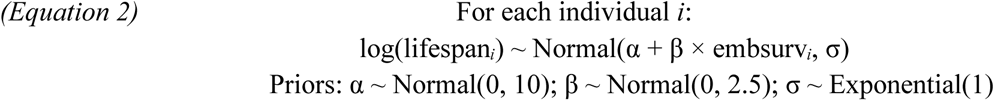

Where: log(lifespan*ᵢ*) = log-transformed lifespan of individual *i;* embsurv = proportion of embryos survived for individual *i;* α= intercept; β= slope (effect of embryo survival on log-lifespan); σ= residual standard deviation. We performed similar models for fertilization rate by using the proportion of fertilized eggs. We assessed model convergence as before (see associated code). As before posterior probabilities of the intercept and slope were extracted (“extract” function) for each age class, and the posterior predictions of lifespan were computed across a regular sequence of 100 embryo survival values. As before, we plotted individual posterior predictions and the mean posterior prediction for each age class using “ggplot2” (see associated code). The same statistical procedures were also applied to fertilization rate (**Supplementary Results**).

### Simulations based on the DAG

To test whether the hypothesized causal model (DAG, **Figure 2a**) could reproduce the observed patterns between lifespan and embryo survival, we used simulated data. We generated a population of 1000 individuals, each assigned an *intrinsic quality* (IQ) produced from a normal distribution (mean = 0, sd = 1). Lifespan (LS) was then simulated for each individual as a log-normal transformation of QI, while functional quality (FQ) was modelled to decline linearly as age increases. At each age (i.e. from 1 until the end of the individual’s lifespan), we generated embryo survival (ES) from a normal distribution with mean determined by FQ and age. Age classes were defined according to population survival thresholds, consistent with the experimental data: “young” when ≥85% of the population was alive, “mid-age” between 85% and 60%, and “old” when ≤60% of the population is left.

We ran Bayesian models in Stan by, similarly to *Equation 2*, modelling lifespan independently for each age class as a linear function of Z-transformed embryo survival. As before, each model was run with 4 chains, each consisting of 2000 iterations, of which 1000 for warmup. We assessed model convergence as above. Posterior probabilities of the intercept and slope were extracted (“extract” function) for each age class, and the posterior predictions of lifespan were computed across a regular sequence of 100 embryo survival values. We plotted individual posterior predictions and the mean posterior prediction for each age class using “ggplot2”.

## Data and Code Availability

The datasets and code underlying this study will be made publicly available on GitHub. A permanent DOI will be assigned via Zenodo upon acceptance of the manuscript.

## Acknowledgements

We are thankful to the Fish Facility of the Leibniz Institute on Aging, especially Christin Hahn, Johannes Wilfert, Ronja Baal, Marcel Münnich, and Magno Delmiro Garcia for providing constant support to our work, and Beate Hoppe and Uta Naumann for the support in managing the ethical approval. We thank all the past and current members of the Valenzano lab for their continuous input and support.

## Supplementary Results

### Male fertilization rate decreases with age and does not trade-off with lifespan

To investigate how investment in reproduction affects lifespan, we performed in vitro fertilizations (IVFs) at three different time points throughout male’s lifespan. As a measure of reproductive success we recorded fertilization rate and embryo survival. Specifically, after 3.5 hours post IVF we checked the fertilization rate by counting the fertilized and unfertilized eggs under a stereomicroscope. The survival of the embryos was then scored 72 hours post fertilization (hpf) by counting the number of embryos still alive. Here we present the results for fertilization rate (i.e. the number of fertilized eggs on the total number of eggs).

In line with previous work, we found that age significantly affected fertilization rate (*X*^2^=56.5612, *p*=5.223e-13). Moreover, sperm concentration significantly affected fertilization rate: males producing more sperm fertilize more eggs (*X*^2^=7.0762, *p*=0.0078). Old males fertilized significantly less eggs than young males (*z*=7.167, *p*<0.0001) (**Figure S1a**). Similarly to embryo survival, fertilization rate, significantly declines already at mid-age (*z*=4.919, *p*<0.0001) (**Figure S1a**).

To test for reproduction-survival trade-offs, we estimated posterior predictive distributions of the relationship between lifespan and fertilization rate at the three different ages (**Materials and Methods**). We found no negative correlation between fertilization rate and individual lifespan (**Figure S1b**). Posterior estimates indicate that males achieving higher fertilization rate at young age live longer (β_mean_=0.906; 0.48 – 1.31, 95% credible intervals), while there is no correlation between individual survival and fertilization rate when sampling males at mid (β_mean_=0.14; -0.38 – 0.31, 95% credible intervals) and old age (β_mean_=-0.61; -1.98 – 0.84, 95% credible intervals) (**Figure S1c**).

### Early fertilization rate predicts individual lifespan

Since father survival and fertilization rate covary, despite weakly, for data collected at young father age (**Figure S1b**), we asked whether fertilization rate statistically predicts variation in father lifespan across individuals. Running a Bayesian model for each age class with father lifespan as the outcome variable (**Materials and Methods, Figure S1d**), we found – as expected from the previous model – that the effect of early-life fertilization rate on lifespan is positive (**Figure S1d**). Specifically, the posterior predictive distribution of the slope at young age (β_mean_=0.34; 0.006 – 0.69, 95% credible intervals) indicates a 95% probability that higher fertilization rate is associated with increased male lifespan (**Figure S1e**). On the contrary, running a model using fertilization rate measured at older ages, the credible intervals associated to the slope of fertilization rate on lifespan include zero both at mid (β_mean_=0.037; -0.49 – 0.52, 95% credible intervals) and old age (β_mean_=-0.13; -0.60 – 0.35, 95% credible intervals), indicating that measuring fertilization rate from mid and old age fathers provides no information about their lifespan (**Figure S1e).**

**Figure S1. Fertilization rate largely mirrors the patterns observed in embryo survival. a)** Boxplot and Kernel density plot depicting fertilization rate at young, mid and old age. All pairwise comparisons are significant. **b)** Scatterplot showing the relationship between lifespan and fertilization rate across the three age classes. Dots represent observed data from each individual male. Mean posterior predictions (black lines) and posterior predictive distributions (coloured lines) from Bayesian models are fitted. For visualization purposes, observed data for fertilization rate and lifespan is transformed using the logit function and the log scale, respectively. **c)** Forest plot of β estimates for each age class. Points indicate posterior means and horizontal bars represent 95% credible intervals, relative to zero as the null effect. **d)** Mean posterior predictions (black lines) and posterior predictive distributions (coloured lines) from Bayesian models run for each age class. Lifespan was modelled as the outcome variable. For visualization purposes, lifespan was back-transformed from the log scale by exponentiation. Black lines represent posterior means. **e)** Forest plot of β estimates for each age class. Points indicate posterior means and horizontal bars represent 95% credible intervals, relative to zero as the null effect.

## Notes

### Competing Interest Statement

The authors have declared no competing interest.

### Summary of Updates

We changed the title and abstract to better reflect the main result, re-ran the Bayesian model, updated Figure 2 and the results section, correspondingly.

### Funding

- Leibniz Institute on Aging - Fritz Lipmann Institute (FLI)

